# Sequence analysis of SARS-CoV-2 genome reveals features important for vaccine design

**DOI:** 10.1101/2020.03.30.016832

**Authors:** Jacob Kames, David D. Holcomb, Ofer Kimchi, Michael DiCuccio, Nobuko Hamasaki-Katagiri, Tony Wang, Anton A. Komar, Aikaterini Alexaki, Chava Kimchi-Sarfaty

**Affiliations:** Center for Biologics Evaluation and Research, Office of Tissues and Advanced Therapies, Division of Plasma Protein Therapeutics, Food and Drug Administration, Silver Spring, MD, USA; Harvard University School of Engineering and Applied Sciences; National Center of Biotechnology Information, National Institutes of Health, Bethesda, MD, USA; Center for Biologics Evaluation and Research, Office of Vaccines Research and Review, Division of Viral Products, Food and Drug Administration, Silver Spring, MD, USA; Center for Gene Regulation in Health and Disease, Cleveland State University, Cleveland, OH, USA

## Abstract

As the SARS-CoV-2 pandemic is rapidly progressing, the need for the development of an effective vaccine is critical. A promising approach for vaccine development is to generate, through codon pair deoptimization, an attenuated virus. This approach carries the advantage that it only requires limited knowledge specific to the virus in question, other than its genome sequence. Therefore, it is well suited for emerging viruses for which we may not have extensive data. We performed comprehensive *in silico* analyses of several features of SARS-CoV-2 genomic sequence (e.g., codon usage, codon pair usage, dinucleotide/junction dinucleotide usage, RNA structure around the frameshift region) in comparison with other members of the coronaviridae family of viruses, the overall human genome, and the transcriptome of specific human tissues such as lung, which are primarily targeted by the virus. Our analysis identified the spike (S) and nucleocapsid (N) proteins as promising targets for deoptimization and suggests a roadmap for SARS-CoV-2 vaccine development, which can be generalizable to other viruses.

## Introduction

The recent emergence of the 2019 novel coronavirus (SARS-CoV-2) has gained worldwide attention and sparked an international effort to develop treatments and a vaccine. To date, there have been 693,224 confirmed cases and 33,106 deaths from COVID-19 worldwide, with 136 countries implementing additional health measures[1]. Given the urgency to combat this emerging disease, multiple efforts to develop an effective vaccine are underway. A relatively recent approach for vaccine development, first proposed by Coleman et al. in 2008 for the attenuation of poliovirus[2], has been used for the attenuation of dozens of viruses, and more recently for bacteria[3]. This approach accomplishes viral attenuation through codon pair deoptimization and appears to be promising for vaccine development, particularly against emerging viruses, as it does not require extensive virus-specific knowledge. It does, however, require knowledge of the viral genome sequence and extensive characterization of its codon and codon pair usage characteristics.

Codon usage is biased across all domains of life, i.e., synonymous codons occur at different frequencies in different organisms[4,5]. It is thought that preferred codons correspond to more abundant tRNAs, and therefore, are translated more efficiently[6]. Similarly, there is bias in codon pair usage, with certain codon pairs occurring at a much different frequency than would be expected based on the codon usage[5]. Codon pair usage also appears to affect translation efficiency[6], although the mechanism is not entirely clear, and it has been argued that dinucleotide usage may be the driving force in determining viral sequence fitness, while codon pair bias may be a secondary effect of altered dinucleotide frequency[7]. Considering that viruses are obligate intracellular parasites and rely on the host-cell machinery for proper expression of their genes, it is worth noting that their codon usage often does not closely resemble the codon usage of their hosts[8,9], a phenomenon that is not well understood. In this regard, a thorough characterization of codon, codon pair and dinucleotide usage of SARS-CoV-2 can provide useful information regarding expression potential of the viral genes and the fitness of the virus in its human or other hosts. Furthermore, it has been shown that viral attenuation can be achieved through extensive changes in codon pair usage of viral genes[2]. Since the mechanism of viral attenuation through codon pair deoptimization is not entirely clear, this in-depth analysis is necessary to guide the development of new vaccines.

Coronaviruses (CoVs) are enveloped, positive-stranded RNA viruses with a large genome of about 30 kb encoding multiple proteins[10]. Translation of a positive-stranded RNA from the initial infectious virus particles generates (among other proteins) a virally encoded RNA dependent RNA polymerase (replicase). This replicase is necessary for viral replication and subsequent generation of viral subgenomic RNAs (sgRNAs), from which the synthesis of structural and accessory proteins occurs[10]. ORF1ab, which encodes the replicase polyprotein (among other proteins) occupies about two thirds of the 5’ prime end of this genome[10,11]. A -1 programmed ribosomal frameshift (PRF) occurs half-way through ORF1ab, allowing the translation of ORF1b[10]. The efficiency of the frameshift thus modulates the relative ratios of proteins encoded by ORF1b and the upstream ORF1a and is critical for coronavirus propagation. Frameshift efficiency (ranging from 15 to 60%) in -1 PRFs is commonly regulated by pseudoknotted mRNA structures following the frameshift, and the conservation of a three-stem pseudoknot in coronaviruses has been previously characterized[12]. Following ORF1ab, are the spike (S), ORF3a, envelope (E), membrane (M), ORF6, ORF7a, ORF7b, ORF8, nucleocapsid (N) and ORF10 genes. S, E, M and N are the structural proteins of the virus[11]. S promotes attachment and fusion to the host cell, during infection[13]. In the case of SARS-CoV-2, S binds to the human angiotensin-converting enzyme 2 (ACE2)[11,14,15]. The E protein is an ion channel and regulates virion assembly[16]. The M protein also participates in virus assembly and in the biosynthesis of new virus particles[17], while the N protein forms the ribonucleoprotein complex with the virus RNA[18] and has several functions, such as enhancing transcription of the viral genome and interacting with the viral membrane protein during virion assembly[19]. Many of the other ORFs have unknown functions or are not well characterized[20], as their presence is not consistent across all coronaviruses.

We have conducted a thorough analysis of the codon, codon pair and dinucleotide usage of the SARS-CoV-2 and have assessed how it relates to other coronaviruses, its hosts, and to the tissues that SARS-CoV-2 has been reported to infect[21-23]. We have taken advantage of our recently published databases, which include genomic codon usage statistics for all species with available sequence data, and transcriptomic codon usage statistics from several human tissues[4,5,24]. We further analyzed each viral gene in terms of its codon characteristics and used an array of codon usage metrics that informed us of the potential of each gene sequence to contribute to the deoptimization of the virus. In the case of ORF1ab, we further examined the structure of the mRNA in the region following the frameshift, finding the SARS-CoV-2 mRNA to exhibit a similar pseudoknotted structure to known coronaviruses. We identified two viral genes that represent valuable targets for deoptimization to generate an attenuated virus. Our analysis can be used as a pipeline to guide codon pair deoptimization for viral attenuation and vaccine development or *a posteriori* to evaluate the effectiveness of an attenuated viral sequence.

## Results

### SARS-CoV-2 proximity to coronaviruses, host genomes and tissue transcriptomes

Since the end of last year when it first emerged, SARS-CoV-2 has been mutating and spreading around the world. Ninety-seven complete or near-complete SARS-CoV-2 genomes are currently accessible in GenBank, with various mutations. To determine which SARS-CoV-2 sequence was most appropriate to use, we retrieved all the published sequences of the virus available in GenBank; after excluding incomplete and low-quality sequences, we calculated the percent difference in codon usage between these and the reference sequence. The average percent difference in codon usage was 0.029%, or ∼3 codons / 10,000, clearly showing that variation in sequences is not significantly affecting overall codon usage. This degree of mutation between strains is corroborated by a recently published study[25].

We next examined how the SARS-CoV-2 codon pair usage compares with other coronaviruses and to its current host. Codon pair data inherently contain the codon usage data and therefore are better suited than codon usage data for this type of comparison. As expected, SARS-CoV-2 codon pair usage closely resembles the codon usage of the coronoviridae family, while it is quite distinct from the codon pair usage of the human genome (Table 1). Bat (*Chiroptera*) and pangolin (*Pholidota*) from which the virus may have been transmitted to humans, as well as dog (*Canis lupus familiaris*) to which the virus is feared may be transmitted next, were included in the analysis. We find that these species have a similar codon usage when compared with human; therefore, viral tropism cannot be inferred based on codon usage data alone (Table 1). Since SARS-CoV-2 infects bronchial epithelial cells and type II pneumocytes and our recent findings show that transcriptomic tissue-specific codon pair usage can vary greatly from genomic codon pair usage[24], we also examined the transcriptomic codon pair usage of the lung and how it compares with the SARS-CoV-2 codon pair usage. Rather surprisingly, the codon usage in the lung was more distinct from SARS-CoV-2 codon usage than the *Homo sapiens* genomic codon usage. The transcriptome codon pair usage of kidney and small intestine, tissues that are also susceptible to the infection, are similarly distant from SARS-CoV-2 (Table 1).

**Table 1.** Euclidean distance (scaled /1000) between codon pair usage frequencies of SARS-CoV-2, Coronaviridae (All CoV), *Homo sapiens* (genomic), lung, kidney (cortex), small intestine (terminal ileum), *Pholidota* (pangolins), *Chiroptera* (bats) and *Canis lupus familiaris*.

|  | SARS-CoV-2 | All CoV | *H. sapiens* Genomic | Lung | Kidney | Small Intestine | Pholidota | Chiroptera |
| --- | --- | --- | --- | --- | --- | --- | --- | --- |
| All CoV | 12.75 |  |  |  |  |  |  |  |
| *H. sapiens* Genomic | 23.20 | 20.79 |  |  |  |  |  |  |
| Lung | 26.18 | 23.52 | 5.80 |  |  |  |  |  |
| Kidney | 26.12 | 23.43 | 6.00 | 1.56 |  |  |  |  |
| Small Intestine | 25.93 | 23.26 | 5.54 | 1.73 | 1.85 |  |  |  |
| Pholidota | 24.69 | 22.16 | 2.76 | 4.18 | 4.46 | 4.09 |  |  |
| Chiroptera | 24.00 | 21.54 | 1.75 | 4.70 | 4.91 | 4.48 | 1.76 |  |
| *Canis lupus familiaris* | 23.68 | 21.24 | 1.28 | 5.18 | 5.37 | 4.92 | 2.18 | 1.04 |

### Codon, codon pair and dinucleotide usage of SARS-CoV-2

To inspect the sequence features of SARS-CoV-2 in more detail, we plotted its codon usage per amino acid and compared it with the human genome and lung transcriptome (Figure 1). SARS-CoV-2 clearly exhibits a preference in codons ending in T and A (71.7%), which is not observed in the human genome (44.9% ending in T or A) and lung transcriptome (37.6% ending in T or A). Similarly, the kidney and small intestine transcriptome show a preference for codons ending in C and G (62.5% in the kidney and 61.8% in the small intestine, Supplemental Figure 1). The codon pair usage of SARS-CoV-2 was also examined in juxtaposition with the human codon usage (Figure 2A and 2B). The differences in codon usage of the two genomes are highlighted in Figure 2C.

**Figure 1.**
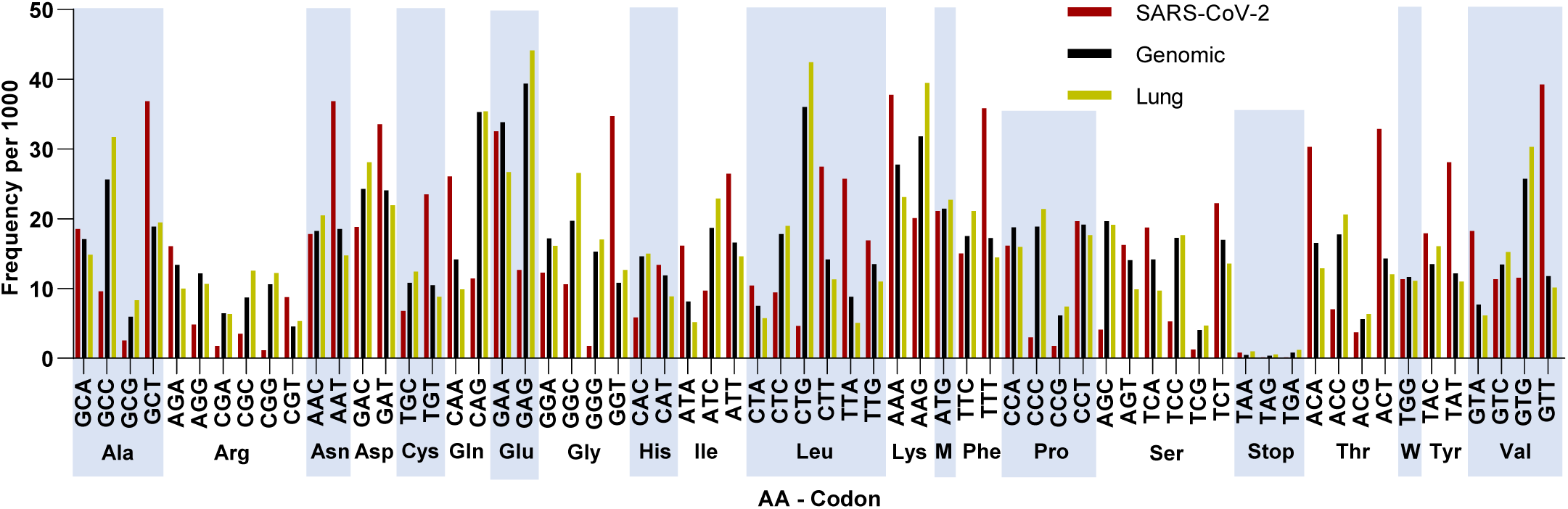
Codon frequencies per 1000 for SARS-CoV-2 (Red), *Homo sapiens* (Genomic, Black) and Lung (Yellow). Codons are grouped by the amino acid they encode (alternating light blue columns, Met and Trp represented as single letter).

**Figure 2.**
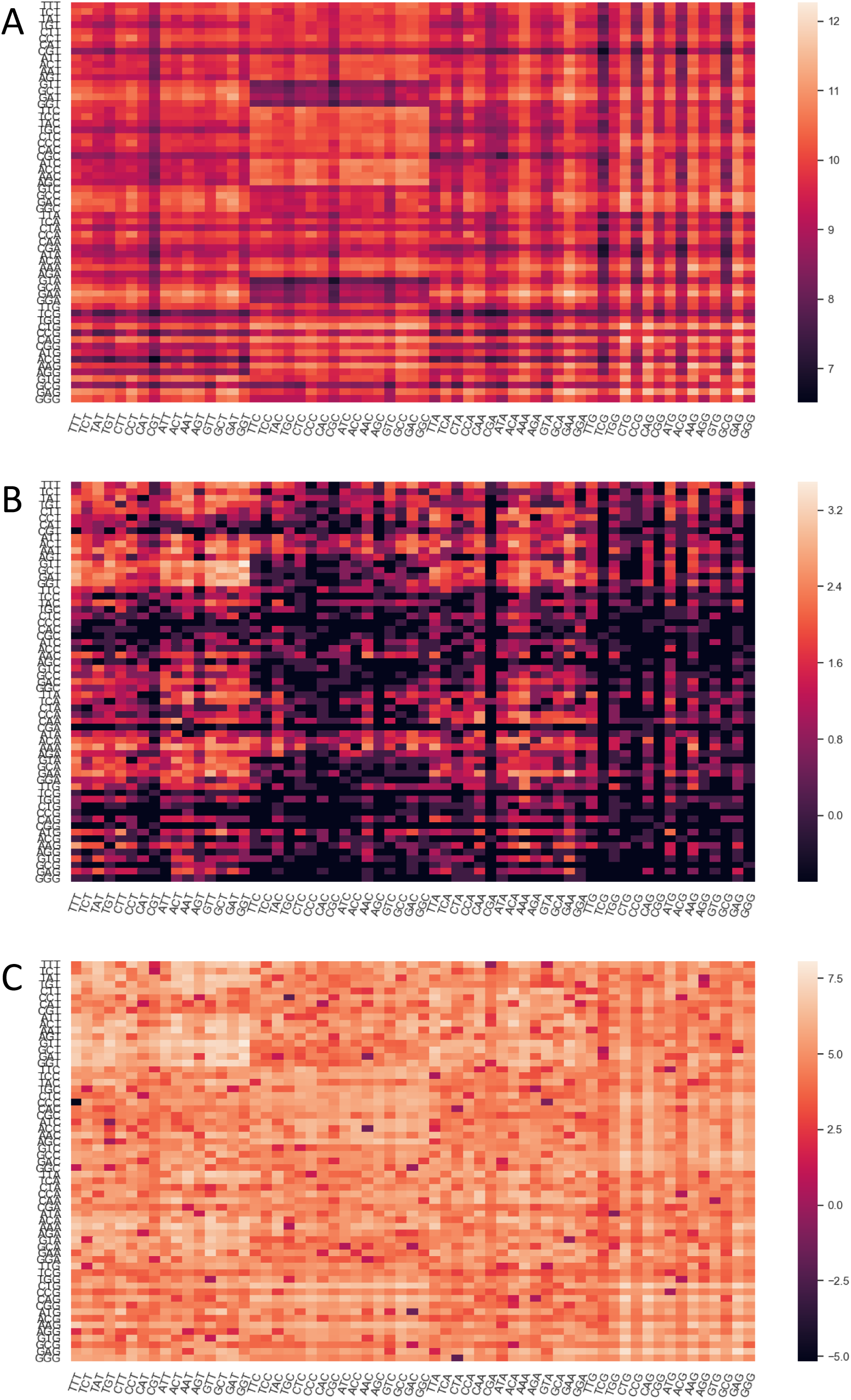
Heat maps of log transformed codon pair frequencies per 1M for *Homo sapiens* (Genomic, **A**), SARS-CoV-2 (**B**) and the absolute value of difference between the two (**C**). Codon pairs increase in frequency from dark to light.

Since the mechanism of viral attenuation through codon pair deoptimization is not entirely clear, and it has been argued that it is an indirect result of increased CpG content, we further investigated the dinucleotide and junction dinucleotide profile of the SARS-CoV-2 as it compares with *Homo sapiens* genome and lung transcriptome (Figure 3). Clearly, CpG dinucleotides are avoided in the SARS-CoV-2 genome, and to a lesser extent CC and GG dinucleotides are too. This provides an opportunity to increase immunogenicity of a potential attenuated virus vaccine by increasing its CpG content.

**Figure 3.**
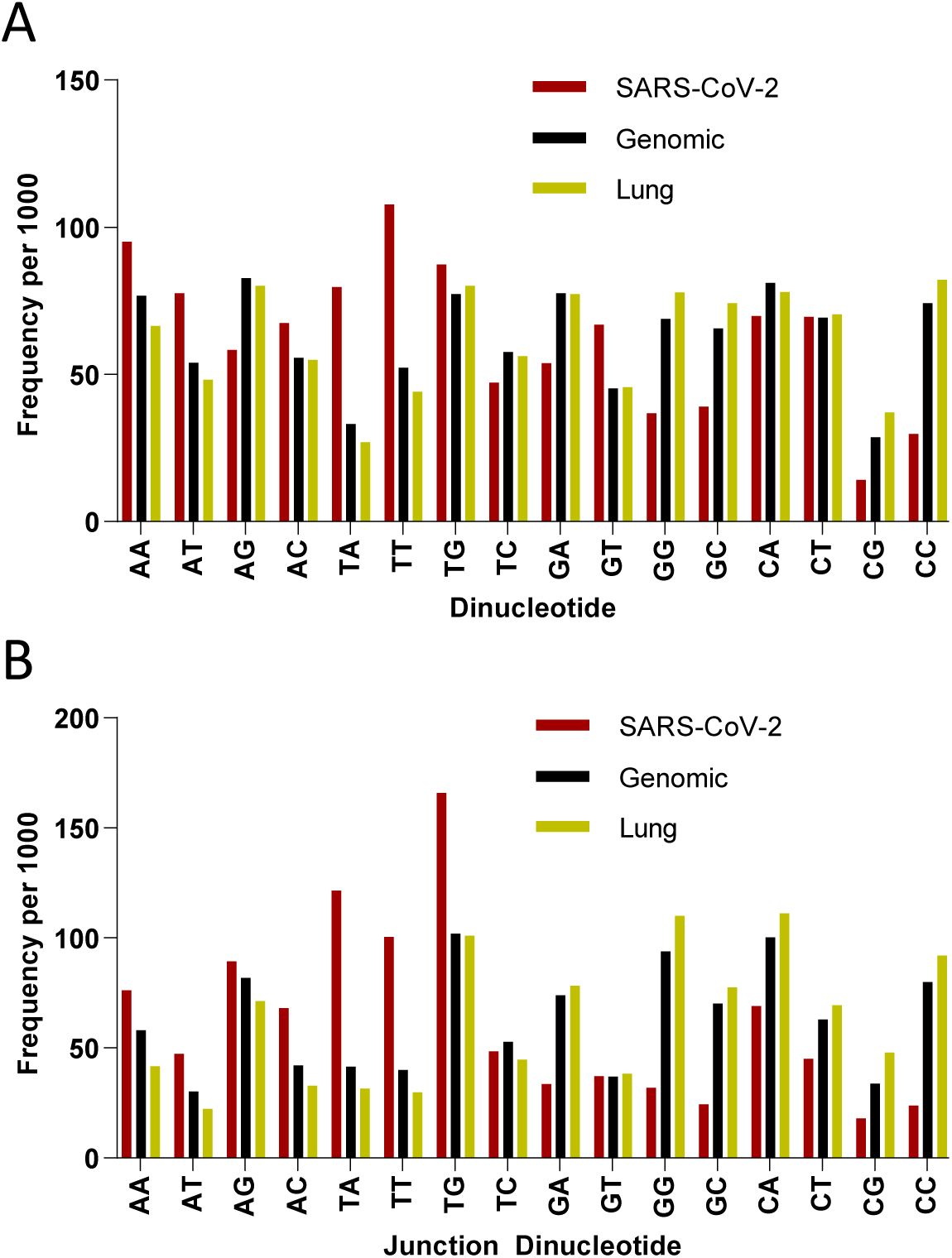
Dinucleotide (**A**) and junction dinucleotide (**B**) frequencies per 1000 for SARS-CoV-2 (Red), *Homo sapiens* (Genomic, Black) and Lung (Yellow).

### RNA folding

The genome sequence determines not only the amino acid sequence, but also the structure of the mRNA. The mRNA structure following the frameshift site is expected to be especially biologically relevant, as pseudoknots following programmed ribosomal frameshifts have been found to regulate the efficiency of the frameshift[26,27]. We therefore sought to study the similarity of the SARS-CoV-2 mRNA structure compared with the structures of different coronavirus mRNAs in the region following the ORF1ab frameshift.

RNA structures were predicted using two distinct secondary structure prediction algorithms[28-30]. Of the top 10 coronaviruses whose predicted minimum free energy (MFE) structures best aligned to that of SARS-CoV-2, seven matched among the two algorithms, showing a high degree of agreement among the two sets of structure predictions. Those seven consensus best-aligned structures are shown, alongside the novel coronavirus post-frameshift structure, in Figure 4 A-H. The similarity of two of these structures to SARS-CoV-2 can be explained by a high degree of sequence similarity to the SARS-CoV-2 mRNA (a SARS-related coronavirus and a bat coronavirus, shown in Figure 4 B-C. However, the other five —all belonging to avian coronaviruses, which are part of the group of the so-called gammacoronaviruses, causing highly contagious diseases of chickens, turkey and other birds —were not in the top 10 sequences most closely aligned to the SARS-CoV-2 mRNA on the basis of sequence. Of note, in the 97 SARS-CoV-2 sequences found in GenBank, no mutations were found in the sequences around the frameshift site (+/- 200 nts) further highlighting the functional importance of this region.

**Figure 4.**
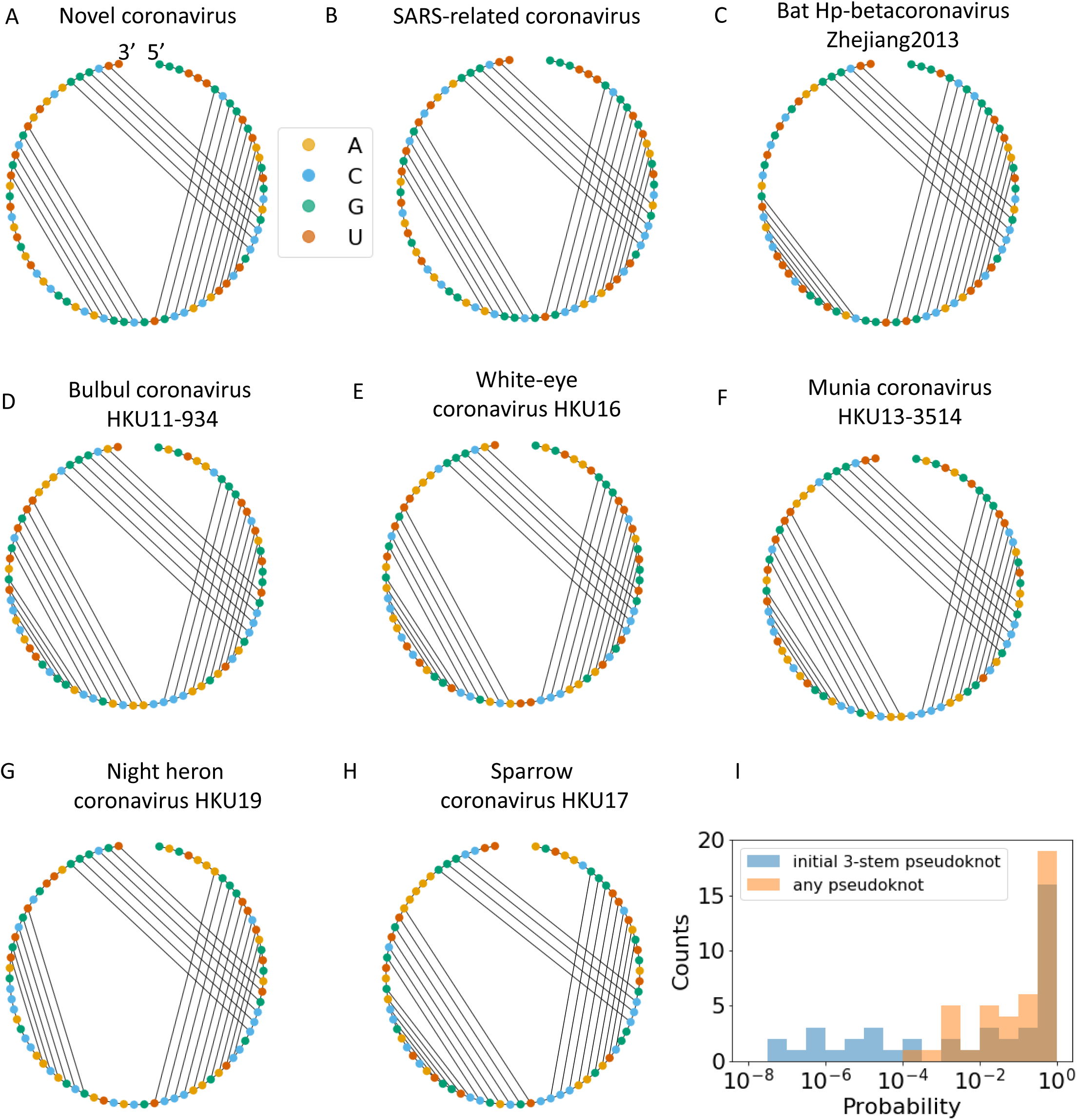
**A**: The predicted minimum free energy (MFE) secondary structure of the novel coronavirus RNA in the 75 nts following the frameshift. All MFE structures displayed are those predicted by our landscape enumeration algorithm; results discussed were found to be insensitive to prediction algorithm by comparison to NuPack. **B-C:** Known coronaviruses with high degree of sequence and structure similarity to the novel coronavirus. **D-H** Known coronaviruses with a high degree of structure similarity to the novel coronavirus, but less sequence similarity. See main text for further discussion. **I:** In addition to examining the predicted MFE structures, we considered the full free-energy landscapes. The probability of each coronavirus to form a pseudoknot in the 75 nts following the frameshift (orange), and the probability of the first stem to be part of a 3-stem pseudoknot (blue), are histogrammed.

Finally, we used our recently published RNA landscape enumeration algorithm[29] to study the RNA folding beyond the MFE structures. We find that even those coronaviruses whose MFE structure does not contain a pseudoknot will fold into a pseudoknot in a relatively high fraction of cases, and that most coronaviruses have a relatively high probability of the initial stem following the frameshift folding into part of a 3-stem pseudoknot like the one exhibited by the SARS-CoV-2 MFE structure (Figure 4 I).

### Viral gene codon usage properties

We next sought to examine each viral gene separately in terms of their codon and codon pair usage. Relative synonymous codon usage (RSCU) and codon pair score (CPS) are commonly used metrics to describe the codon and codon pair usage bias, respectively. RSCU expresses the observed over expected synonymous codon usage ratio, while CPS is the natural log of the observed over expected synonymous codon pair ratio using observed individual codon usage[2,31]. In our analyses, RSCU and CPS are derived from human genomic codon and codon pair usage frequencies. For ease of comparison, we used ln(RSCU) to measure the codon usage bias. The average CPS across a gene is referred to as codon pair bias (CPB) of the gene[2]. The average ln(RSCU) and CPB of each viral gene was calculated and compared with host genes average ln(RSCU) and CPB (Figure 5). The average RSCU, ln(RSCU) and CPB of each viral gene appear in Table 2. ORF10 was strikingly the least similar gene to the human genome in terms of both its codon and codon pair usage, followed by the E gene. These genes provide little opportunity for deoptimization, since their sequence is already far from optimal. On the other hand, genes S and N are more similar to human in terms of their codon pair usage. To explore further the potential for codon pair deoptimization, we plotted their CPS across their sequence (Supplemental Figure 2 and Figure 6). As seen in these figures all viral genes use mostly rare codons (ln(RSCU) <0); however, it is striking that ORF6 and ORF10 use almost exclusively rare codons, while ORF3a and M and ORF10 have some of the lowest ln(RSCU) values. Regarding codon pair usage, S stands out as the gene that uses frequent codon pairs more often (peaks with relatively high CPS scores), while N, ORF6 and ORF7b are genes that do not use very rare codon pairs (CPS values are only moderately negative).

**Table 2.** Codon and codon pair metrics of SARS-CoV-2 genes. Relative synonymous codon usage (RSCU), ln(RSCU) and codon pair bias (CPB) for 11 viral genes.

|  | ORF1ab | S | ORF3a | E | M | ORF6 | ORF7a | ORF7b | ORF8 | N | ORF10 |
| --- | --- | --- | --- | --- | --- | --- | --- | --- | --- | --- | --- |
| Avg RSCU | 0.98 | 1.00 | 0.99 | 0.97 | 1.01 | 0.95 | 1.04 | 1.02 | 0.98 | 1.02 | 0.88 |
| Avg ln(RSCU) | -0.05 | -0.04 | -0.05 | -0.10 | -0.04 | -0.08 | 0.00 | -0.03 | -0.07 | -0.02 | -0.17 |
| CPB | 0.03 | 0.07 | 0.03 | -0.10 | 0.00 | 0.02 | -0.03 | -0.04 | 0.00 | 0.03 | -0.17 |

**Figure 5.**
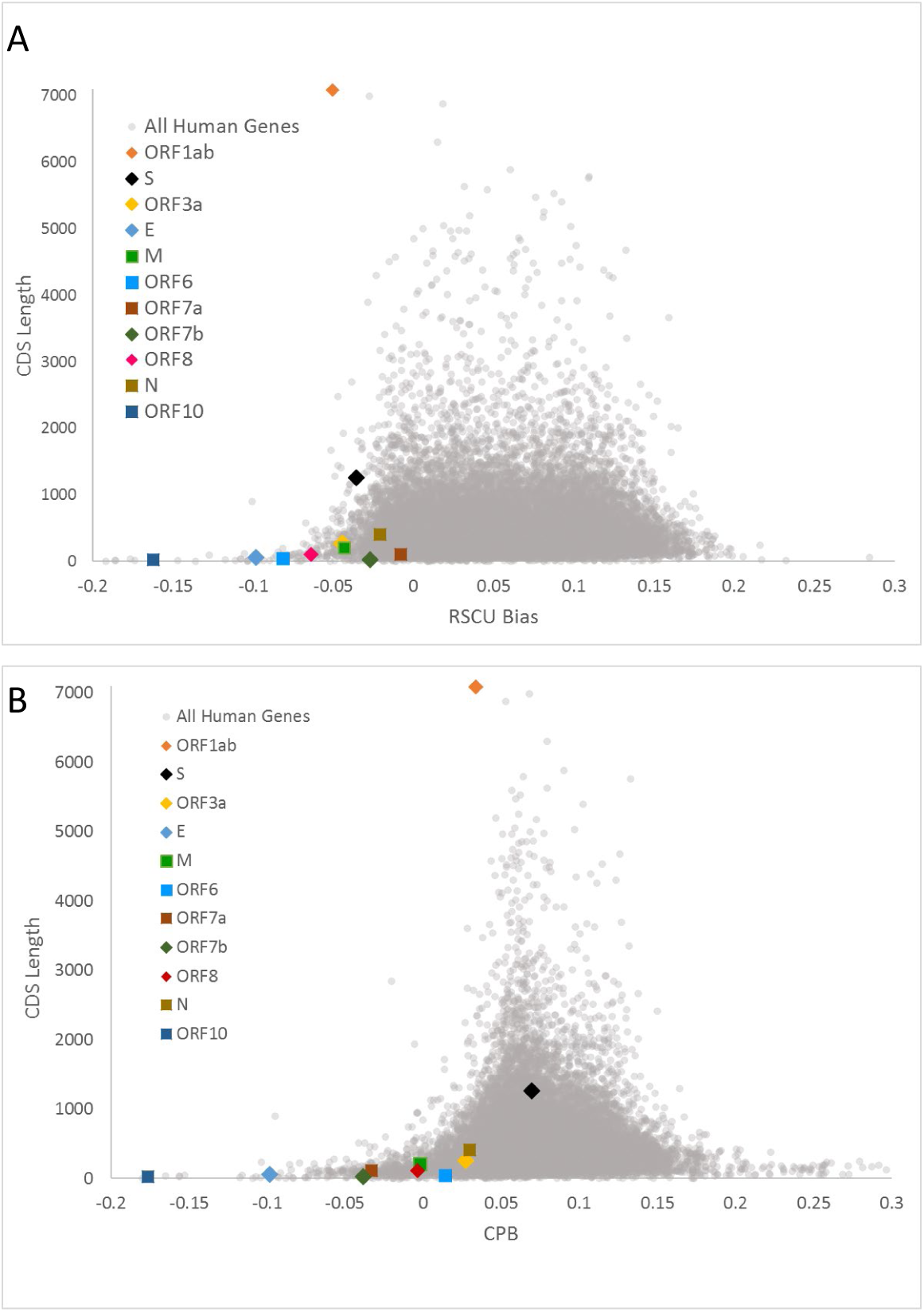
Scatterplots of RSCU bias [average ln(RSCU)] (**A**) and Codon Pair Bias (**B**) by CDS length of human and viral genes. Human genes appear as grey dots and viral genes appear with different colored markers.

**Figure 6.**
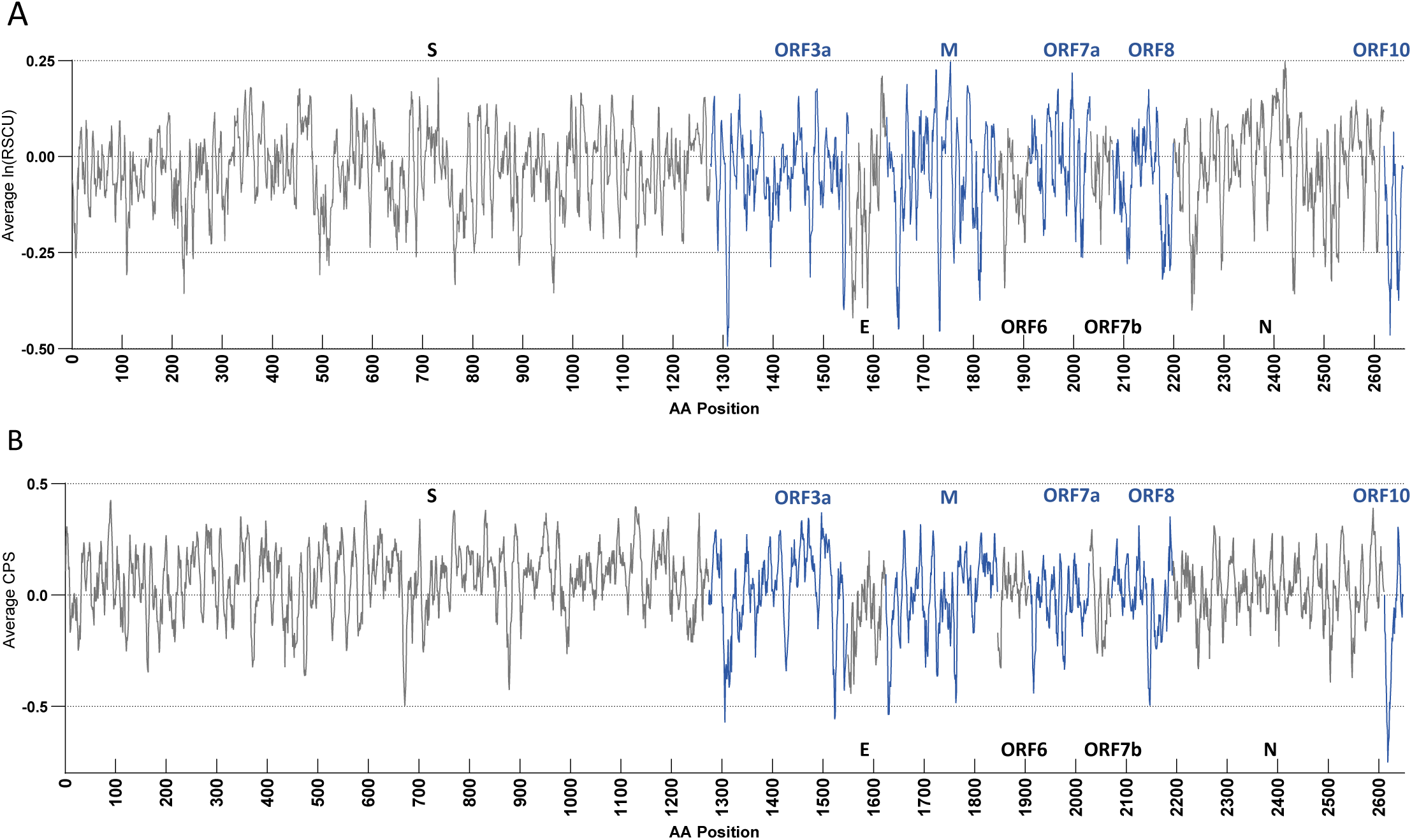
Seven codon sliding window average of ln(RSCU) (**A**) and codon pair score (CPS) (**B**) of structural SARS-CoV-2 genes. Genes are shown in the order they appear in the viral genome, but gaps between open reading frames have been removed. Genes alternate in colors black and blue for clarity, with the gene name in the corresponding color appearing above or below the window. RSCU and CPS are calculated based on *Homo sapiens* genomic codon and codon pair usage.

## Discussion

We performed a comprehensive characterization of the codon, codon pair and dinucleotide usage of the SARS-CoV-2 genome with the intention to identify the best targets for codon pair deoptimization in order to design an attenuated virus for vaccine development. Genes N and S were singled out as the best potential targets for deoptimization, due to the relatively high CPB among the viral genes. Furthermore, they are structural proteins with known functions. Therefore, it will be possible to study the results of codon pair deoptimization on the fitness of the virus.

Currently most published attempts of viral attenuation through codon pair deoptimization do not discuss the strategy for selecting which genes to deoptimize. Although codon pair deoptimization has been proven successful for viral attenuation[2,32,33], the mechanism is not clear. In addition, it is reasonable to assume that for every successful published deoptimization attempt, there may be several unsuccessful and therefore unpublished ones. Similarly, there have been successful attempts to generate attenuated viruses through codon deoptimization[34-37]. Understanding the mechanism that leads to viral attenuation requires a thorough characterization of the viral sequence and of the consequences of sequence changes. There are several factors that may contribute to the efficacy of deoptimization strategies. In changing the codon pair usage, the dinucleotide frequency and the GC content are altered; mRNA secondary structure and translational kinetics are also perturbed. Further, the CpG content is changing, leading potentially to altered immunogenicity. It is likely that codon pair (or codon) deoptimization leads to reduced expression, either due to changes in transcription, mRNA stability, or translation efficiency[38]. Alternatively, it is possible that deoptimization may lead to perturbed cotranslational folding[39], resulting in altered protein conformation. In the case of the S protein, this may lead to decreased binding affinity for the ACE2 protein, thus affecting viral fitness.

A number of parameters were considered in determining which proteins could be targets for codon pair deoptimization. It has been shown that deoptimizing one third or less of the virus is sufficient to attenuate the virus, and more extensive deoptimization may lead to a completely inactive virus[40]. ORF1ab takes up about two thirds of the virus; therefore, its size may make it an unsuitable target. Furthermore, altering its RNA sequence is likely to disturb the pseudoknot that is responsible for frameshifting. Since we have identified the sequence that is responsible for the frameshift, a partial codon pair deoptimization is possible. However, ORF1ab is essential for genome replication, which also does not support its capacity as a codon deoptimization target.

ORF10 has strikingly low CPB and RSCU given that it is at the very end of the viral genome there may be a structural reason for its nucleotide sequence. The E gene also has a very low CPB; interestingly, although ORF10 has both positive and negative CPS across its sequence, E has mostly negative CPSs, which make it unsuited for codon pair deoptimization. ORF7a is unusual, as it has the highest RSCU of the viral genes, but a rather low CPB (it uses preferred codons in unusual combinations). Although ORF7a is not the most compelling target for codon pair deoptimization, if a codon deoptimization strategy is attempted, this gene should be considered. It should, however, be noted that since it overlaps for a few nucleotides with ORF7b, any sequence changes should be considered in coordination for both genes.

Our sequence analysis pointed to the S and N genes as potential targets for codon pair deoptimization. The S protein, which binds to the cell-surface receptor and induces virus-cell membrane fusion, has the highest CPB score of all viral proteins, leading to significant flexibility for codon pair deoptimization. Since it is a surface protein, it is under constant pressure to avoid the immune response, and as a result, it is the least conserved of the coronavirus proteins, diverging both in its amino acid and its codon usage[25]. Being under constant selective pressure could be the reason why it has been able to adapt to the host genome more than other proteins. The N protein forms the ribonucleoprotein complex with the virus RNA. In contrast to S, the N protein is the most conserved and stable protein among the coronavirus structural proteins. It uses mostly codon pairs with intermediate frequency; thus, it could be substantially codon pair deoptimized.

Our thorough codon/codon pair characterization of the SARS-CoV-2 genome has led to the identification of S and N as potential targets for codon pair deoptimization. The next obvious step is to construct the deoptimized virus and test its infectivity and ability to replicate. While testing the fitness of the virus, the strategy of selecting targets would also be tested, which could lead to better understanding of the factors that make codon pair deoptimization successful in generating attenuated viruses. The risk of a new emerging virus is always present, and this has been poignantly highlighted by the current SARS-CoV-2. The current work could be used for the quick generation of a SARS-CoV-2 vaccine but also as a pipeline to facilitate vaccine development when the next virus is presented.

## Materials and Methods

### Sequence Accession and Codon Comparison

The complete reference sequences for severe acute respiratory syndrome coronavirus 2 (SARS-CoV-2, accession NC_045512.2) was downloaded from NCBI RefSeq[41] on March 13, 2020. CDS sequences from 97 complete SARS-CoV-2 isolates were downloaded using NCBI Batch Entrez on March 25, 2020. Sequences of poor quality or with CDS lengths that did not match those of the reference sequence due to deletion or insertion were removed, leaving 87 sequences. To calculate percent difference in codon usage, each CDS of the 87 sequences was compared at the codon level to that of the reference sequence. Codons containing nucleotides where a base call could not be made (“N”) were removed from the calculation. All scripts for this calculation were written in Python 3.7.4.

### Comparison of Codon and Codon Pair Usage in Host Species

Codon, codon pair and dinucleotide usage data for *Homo sapiens, Canis lupus familiaris, Chiroptera* (bats) and *Pholidota* (pangolins) were downloaded from the CoCoPUTs database[5] on March 13, 2020. Likewise, human lung, kidney (cortex) and small intestine (terminal ileum) tissue-specific codon, codon pair and dinucleotide usage data were accessed from the TissueCoCoPUTs database[24] on March 13, 2020. Codon, codon pair and dinucleotide usage data for SARS-CoV-2 was calculated from the reference sequence (accession NC_045512.2) using scripts written in Python 3.7.4. Euclidean distances between codon pair usage frequencies were calculated using the *dist* function from the *stats* package in R 3.6.1.

### RSCU, CPS and CPB

RSCU was calculated as defined in Sharp *et al*.[31] based on *Homo sapiens* genomic codon usage data accessed from the CoCoPUTs database[5] on March 13, 2020. Codon pair scores (CPS) for all 4096 codon pairs were calculated as described in Coleman *et al*[2] using *Homo sapiens* genomic codon pair usage data accessed from the CoCoPUTs database[5] on March 13, 2020. Codon pair bias (CPB) of a gene is the arithmetic mean of all CPSs throughout the gene, as defined in Coleman *et al[2]*.

### RNA folding

To ensure our results are robust to the prediction algorithm chosen as well as to the size of the window examined, we used two secondary structure prediction algorithms on two window sizes. We used NuPack to predict the minimum free energy (MFE) secondary structure on the 100 nucleotides (nts) following the frameshift, and our own recently published free energy landscape enumeration algorithm to examine the full structure landscape of the 75 nts following the frameshift[28,29]. For the latter, we employed the heuristic that the minimum stem length was set to 4. Aside from this heuristic, the two algorithms differ primarily in the loop entropy calculation, which especially affects the probability of pseudoknot formation.

Sequence alignment was measured using MatLab’s Needleman-Wunsch sequence alignment implemented on the 100 nts following the frameshift using default parameters. The parameters employed were the defaults: the NUC44 scoring matrix and a gap penalty of 8 for all gaps.

Structure alignment was measured using a method similar to our previously-studied “per-base topology” score[29]. Taking the dot-bracket representation of each secondary structure, we summed the number of positions containing identical elements. Employing an alignment model allowing for gaps, with a gap penalty and a misalignment penalty of -1, did not change our results[42].

## Author contributions statement

J.K., D.D.H. and O.K. conducted analysis, verified analytic methods, prepared figures and assisted in preparing the original manuscript.

M.D., N.H-K., T.W. and A.A.K. suggested analyses, conducted critical review of the data and assisted in preparing the original manuscript.

A.A. and C.K-S. conceived the original idea, suggested analyses, conducted critical review of the data and prepared the original manuscript.

## Additional information

### Funding

This work was supported by funds from the U.S. Food and Drug Administration Chief Scientist grant and in part supported by the National Institutes of Health grant HL121779 (A.A.K.).

### Conflict of Interest

The authors declare no competing interests.

