## Supplemental Figure for "Sequence analysis of SARS-CoV-2 genome reveals features important for vaccine design"

### Supplemental Figure 1

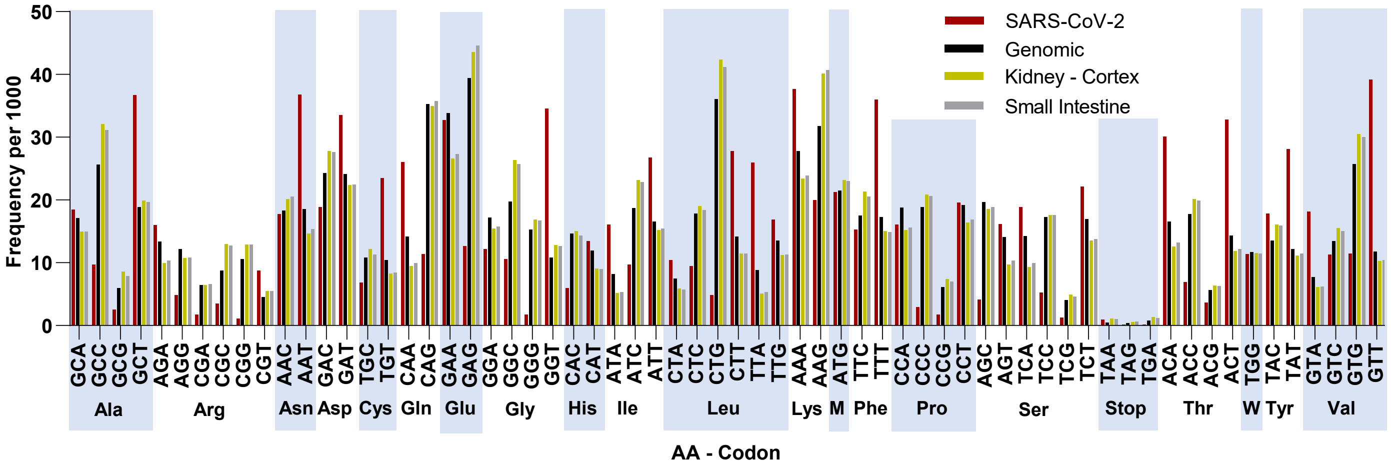


**Supplemental Figure 1** – Codon frequencies per 1000 for SARS-CoV-2 (Red), *Homo sapiens* (Genomic, Black), Kidney (Cortex, Yellow) and Small Intestine (Terminal Ileum, grey). Codons are grouped by the amino acid they encode (alternating light blue columns, Met and Trp represented as single letter).

### Supplemental Figure 2

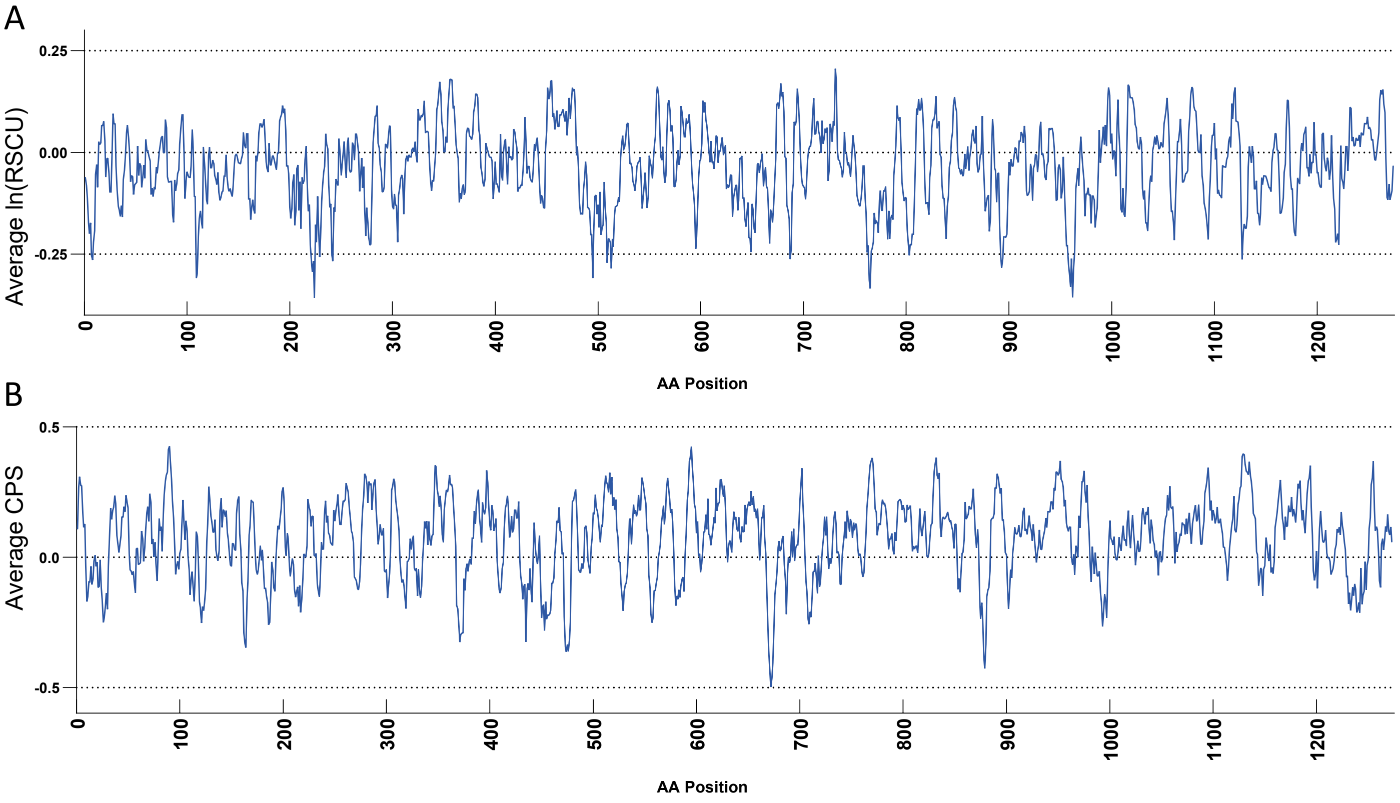


**Supplemental Figure 2** – Seven codon sliding window average of ln(RSCU) (**A**) and codon pair score (CPS) (**B**) of SARS-CoV-2 ORF1ab. RSCU and CPS are calculated based on *Homo sapiens* genomic codon and codon pair usage.
